# Large-scale brain network differences during deliberate and spontaneous mind-wandering in a sustained attention task: An electroencephalography source-level connectivity study

**DOI:** 10.64898/2026.01.22.700987

**Authors:** Xiang Yan, Toru Takahashi, Yuto Kurihara, Masato Ito, Rieko Osu

## Abstract

Mind-wandering (MW), the shift of attention away from an ongoing task, can be classified as either deliberate or spontaneous, depending on whether these internal thoughts arise intentionally or unintentionally. Previous functional magnetic resonance imaging-based resting state studies have shown that spontaneous MW traits are positively correlated with coupling within the default mode network (DMN), whereas deliberate MW traits are positively correlated with coupling both within the DMN and between the DMN and control-or attention-related networks. However, state-based functional connectivity during each MW episode remains unknown. Addressing this knowledge gap, we investigated how deliberate and spontaneous MW differentially involve state-based large-scale network interactions during a sustained attention task using electroencephalography (EEG), which captures rapid network dynamics. Thirty-one participants performed the gradual-onset continuous performance task using 64-channel EEG. Mental states were classified via experience-sampling probes as on-task, deliberate, or spontaneous MW. EEG data from 1-s pre-probe epochs were analyzed using source estimation and the weighted phase lag index to assess functional connectivity within and across the DMN, control network (CN), dorsal attention network (DAN), and salience network (SN) in the delta, theta, alpha, and beta frequency bands. Relative to both spontaneous MW and the on-task state, deliberate MW was characterized by significantly stronger alpha band functional connectivity. A prominent connectivity cluster was centered on the right frontal operculum–insula of the SN, linking regions across the SN, CN, and DMN. In addition, deliberate MW exhibited enhanced SN−DAN−DMN coupling compared with the on-task state. By contrast, spontaneous MW did not show significant connectivity differences relative to the on-task state in any frequency band. Taken together, these results suggest that alpha band network interactions centered on the SN may contribute to the intentional regulation of internally directed thought during deliberate MW.

## 1. Introduction

Mind-wandering (MW) is a type of internal thinking that occurs when an individual performs a task, but the thinking content is not directly related to the task itself (Smallwood & Schooler, 2006). While MW has been associated with reduced cognitive control (Mrazek et al., 2012; Unsworth & McMillan, 2013), other evidence also suggests that individuals with stronger cognitive control capability tend to experience it more frequently during easy tasks (Bernhardt et al., 2014; Rummel & Boywitt, 2014; Smallwood et al., 2013). This indicates that cognitive control may not only suppress MW but may sometimes facilitate the construction and regulation of task-irrelevant internally directed thoughts, as proposed in previous studies (e.g., Christoff et al., 2016; Kucyi et al., 2023). A critical factor underlying this duality may be whether top-down control systems intentionally regulate internal cognition, giving rise to deliberate or spontaneous forms of MW (Golchert et al., 2017; Seli, Carriere, et al., 2015).

The spontaneous MW trait refers to the tendency to have unintentional, drifting thoughts in everyday life (Seli, Carriere, et al., 2015) and has been linked to small working memory capacity and attentional lapses (Robison & Unsworth, 2018). The default mode network (DMN), typically characterized as a task-negative network (Greicius et al., 2003; Raichle et al., 2001), supports internally guided processes such as remembering and envisioning the future that are commonly observed during spontaneous thought (Andrews-Hanna et al., 2010, 2014). The spontaneous MW trait has commonly been associated with stronger DMN activity and coupling based on resting-state functional magnetic resonance imaging (fMRI) evidence ( Golchert et al., 2017; Orwig et al., 2023), reflecting the DMN’s internally oriented nature. A higher tendency of spontaneous MW, but not deliberate MW, is also associated with attention deficit hyperactivity disorder (ADHD) (Arabacı & Parris, 2018; Seli, Smallwood, et al., 2015), which is characterized by impairments in attentional regulation and inhibitory control (Bozhilova et al., 2018; Gao et al., 2019). In addition, evidence from previous studies has demonstrated that individuals with higher ADHD scores show reduced functional connectivity between the control network (CN) and ventral visual cortex, along with less detailed internal thought, which together suggest that they may have weakened control over ongoing cognition during tasks (Vatansever et al., 2019). Thus, these findings suggest that spontaneous MW may be associated with the dysregulation of control systems.

By contrast, deliberate MW, characterized by a conscious shift of cognitive focus inward (Seli, Carriere, et al., 2015) and often accompanied by low motivation for the external task (Robison & Unsworth, 2018), has been theorized to rely on executive control processes that help control and organize internally directed thoughts (e.g., Christoff et al. 2016). A higher tendency to engage in deliberate MW has been associated with greater resting-state functional connectivity between the DMN and CN (Golchert et al., 2017), a network that plays a key role in executive control and the intentional guidance of goal-directed behavior (Dixon et al., 2018; Smallwood et al., 2012). Accordingly, divergent neural network associations observed at the trait level may extend to state-level differences between deliberate and spontaneous MW episodes. While previous research has examined the neural correlation of deliberate and spontaneous MW traits (Golchert et al., 2017), studies investigating neural networks of state MW during task performance have not differentiated between the two forms (e.g., Denkova et al., 2019; Groot et al., 2021; Kucyi et al., 2021; Maillet et al., 2019; Mooneyham et al., 2017). Consequently, direct neural evidence distinguishing the two forms of MW during their occurrence in task contexts remains scarce.

In this study, we investigated the differences in large-scale network interactions between deliberate MW, spontaneous MW, and on-task focus using electroencephalography (EEG) source-level connectivity, which captures rapid network dynamics (Hassan & Wendling, 2018). Using online experience sampling (thought-probe) during a sustained attention task, we classified EEG data into three states (deliberate MW, spontaneous MW, and on-task) and applied source estimation to assess functional connectivity across major brain networks. Based on previous trait-level findings (e.g., Golchert et al., 2017; Orwig et al., 2023) and theoretical models (e.g., (Bozhilova et al., 2018; Christoff et al., 2016; K. C. R. Fox et al., 2015), we hypothesized that deliberate MW, compared with both spontaneous MW and on-task states, would be marked by enhanced coupling between the DMN and networks implicated in cognitive control and salience processing. Second, we hypothesized that spontaneous MW, relative to the on-task state, would be marked by increased connectivity within the DMN, and weaker engagement of control- and attention-related network interactions. Finally, we explored whether functional connectivity (FC) patterns differentiating deliberate and spontaneous MW would be differentially associated with attentional tendencies, including ADHD-related traits. We hypothesized that FC patterns differentiating deliberate from spontaneous MW would relate differently to ADHD tendencies: FC during spontaneous MW would be negatively associated with ADHD tendencies, while FC during deliberate MW would not, consistent with evidence that ADHD tendencies relate to spontaneous, not deliberate, MW (Arabacı & Parris, 2018; Seli et al., 2015).

## 2. Methods

### 2.1 Participants

A total of 44 participants without a diagnosis of neurodevelopmental, mood, or anxiety disorders (male = 23; female = 21; age = 21.6 ± 2.5) were recruited for this study. Participants were instructed to obtain at least 7 h of sleep the night before the experiment and abstain from medications (for 24 h), caffeine (for 6 h), and alcohol (for 6 h) prior to the EEG recording. These instructions were sent via email one day before the experiment, and compliance was confirmed on the day of the session. The study was approved by the Office of Research Ethics at Waseda University (2019-153), and written informed consent was obtained from all participants.

### 2.2 Questionnaires

All participants were asked to complete self-reported questionnaires. The Japanese version of the Conners’ Adult ADHD Rating Scale (CAARS) (Macey, 2003) was used to measure ADHD tendencies. The CAARS was measured on a 4-point scale. The subscales of inattention/memory (12 questions), hyperactivity/restlessness (12 questions), impulsivity/emotional lability (12 questions), and the ADHD index were used for the analyses. We used the Mind-Wandering Scale (MWS) (Carriere et al., 2013), translated into Japanese (Yamaoka & Yukawa, 2019), to measure MW traits. It consists of two subscales: deliberate MW (MWS-D) and spontaneous MW (MWS-S). The MWS was measured on a 7-point scale, and each subscale contained four questions.

### 2.3 Experiment paradigm

#### 2.3.1 Sustained attention task

The participants were then asked to perform a gradual-onset continuous performance task (gradCPT) (Esterman et al., 2013; Kucyi et al., 2016), which included thought probes. The task consisted of a set of 10 city and 10 mountain scenes that gradually transitioned from one scene to another, with each transition lasting approximately 1 s. Randomly selected scenes were presented in each trial of the study. During the task, participants were instructed to use their right index finger to respond to target stimuli: city scenes (go trials; 90%), which required a button press response, and mountain scenes (no-go trials; 10%), which required withholding a response (Kucyi et al., 2016). A scrambled image presented before and after the thought probe was used as a non-target stimulus. The final scene before the thought probe switched to a scrambled image was displayed without gradually fading out. After the thought-probe period, a scrambled image was fully displayed without a fade-in, and then gradually faded out into the next target stimulus (city or mountain image).

The participants were asked to answer a series of thought-probe questions based on their thoughts. The thought probe was presented randomly every 30–60 s throughout the task, and in total, 27 thought probes were presented. Each thought probe question was presented with a 20-s response limit. If the participant did not respond within this period, the next probe was displayed. Participants who failed to respond to two consecutive thought probes triggered automatic task termination and were excluded from the analyses.

During the task, participants completed a series of thought-probe questions presented on the screen. Responses were recorded as integer values from 0 to 100 on a visual analog scale, with the numerical values not being visible to the participants. The first question (Q1) assessed task focus: *“To what degree was your focus just on the task or on something else?”* (0 = only else, 100 = only task). The second question (Q2) evaluated metacognitive awareness: *“To what degree were you aware of where your focus was?”* (0 = unaware; 100 = aware). If participants reported an off-task focus (Q1 ≤ 50), they were asked an additional question (Q3): *“Which of the following best characterizes your mental state before the thought probe?”* with two response options: *deliberate* or *spontaneous MW*. Finally, participants rated their sleepiness (Q4) using the 9-point Karolinska Sleepiness Scale (KSS; 1 = very alert, 9 = very sleepy, fighting sleep) (Shahid et al., 2012). Data from Q2 and Q4 were not included in the final analysis.

Participants were provided written instructions along with two practice sets, both of which included two thought probes, to ensure that they understood how to perform the gradCPT and respond appropriately to the thought probes. The task instructions are provided in the Supplementary Materials.

#### 2.3.2 Behavioral data analysis

The reaction time (RT) was determined by assigning a response to each scene transition based on the time elapsed since the start of the current scene (i.e., 0%). Responses to the city image were considered correct if they were based solely on the current image and were not influenced by the preceding or following images. For example, an RT of 900 ms would occur when the button was pressed when the scene was 90% coherent, and the previous scene was 10% coherent. For rare trials with deviant fast (<70% coherence for the current scene) or slow (>40% coherence for the next scene) responses or multiple buttons, RT was calculated using a prior research method explained further below (Esterman et al., 2013; Kucyi et al., 2016).

Ambiguous responses that might indicate a response to the preceding or following trial were assigned to one of the two trials if that trial had no response. In addition, an ambiguous response was assigned to the nearest trial if both adjacent trials had no response, unless one of the trials was a no-go target. Additionally, if a trial included multiple responses, the fastest response was assigned to calculate the RT. Following this procedure, the RT was calculated and classified as a correct response or an error.

### 2.4 EEG recording

EEGs were recorded using the ActiveTwo system (BioSemi B.V., Amsterdam, Netherlands) with 64 electrodes arranged according to the 10–20 system. Electrooculography was simultaneously recorded but the data were not included in the analyses. The sampling rate was set to 2,048 Hz, and the electrode impedance was maintained below 5 kΩ. EEG data were recorded during two 5-min resting-state sessions (eyes closed and eyes open), followed by an approximately 45-min gradCPT task. EEG data from the gradCPT task were analyzed.

### 2.5 EEG processing

EEG recordings were processed offline using MATLAB and the Statistics Toolbox software (MathWorks, Inc., Natick, Massachusetts, United States). The EEG data were first down-sampled to 500 Hz. A high-pass filter with a cutoff of 1 Hz was applied, followed by a 50 Hz notch filter to remove power line noise, and a low-pass filter with a cutoff of 45 Hz was applied to complete the bandpass filtering. The EEGLAB toolbox v 2021.1 (Delorme & Makeig, 2004), which has the function clean_raw_data (Kothe & Makeig, 2013), was used for artifact subspace reconstruction(ASR)-based artifact correction and channel rejection. This was followed by spherical interpolation for channel interpolation. Re-referencing was estimated on average for all EEG channels. Independent component analysis (ICA) was performed to identify and remove artifact components related to eye movements, muscle activity, channel noise, and line noise, using the ICLabel plugin (Pion-Tonachini et al., 2019), and components with more than 70% probability of being artifacts were rejected.

### 2.6 Source-level EEG analysis

#### 2.6.1 Source estimation

The source activity was assessed using the Brainstorm toolbox, in which the minimum norm imaging CBM152 MRI brain template and deep brain structures were used for source estimation. A three-layer boundary element model, generated using the OpenMEEG (Gramfort et al., 2010) project software (http://www.biomedical-engineering-online.com/content/9/1/45), was then utilized to construct the lead field matrix. Source activity was estimated using minimum norm imaging (Baillet et al., 2001) and further refined using sLORETA (Pascual-Marqui, 2002). Subsequently, the orientation-unconstrained source model, which contains three time series of one voxel, was computed and flattened to one time series through principal component analysis (PCA). The ROIs were determined based on Schaefer’s model (Schaefer et al., 2018), which comprised 64 regions of interest (ROI) from the DMN (24 ROIs), CN (13 ROIs), dorsal attention network (DAN) (15 ROIs), and salience network (SN) (12 ROIs) (Table 1).

**Table 1.**
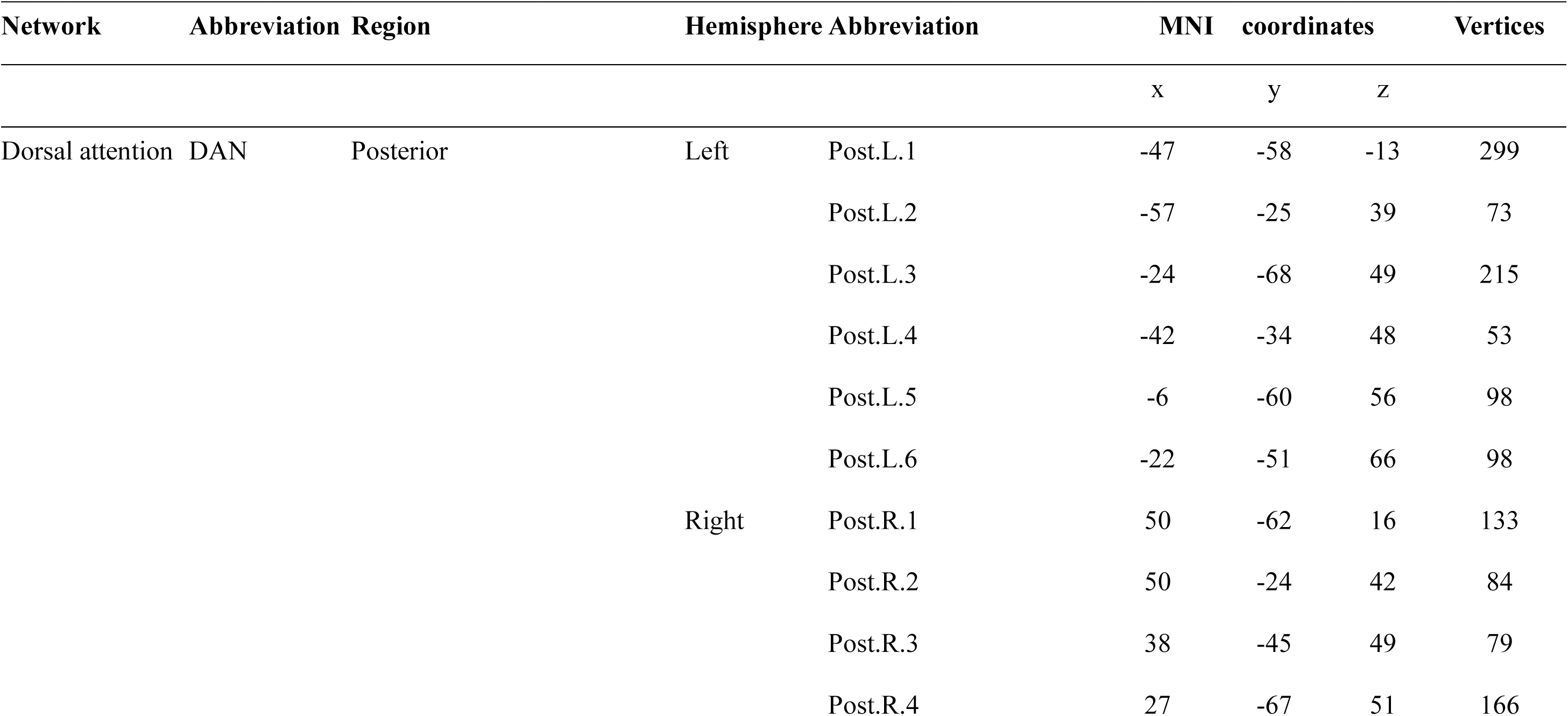
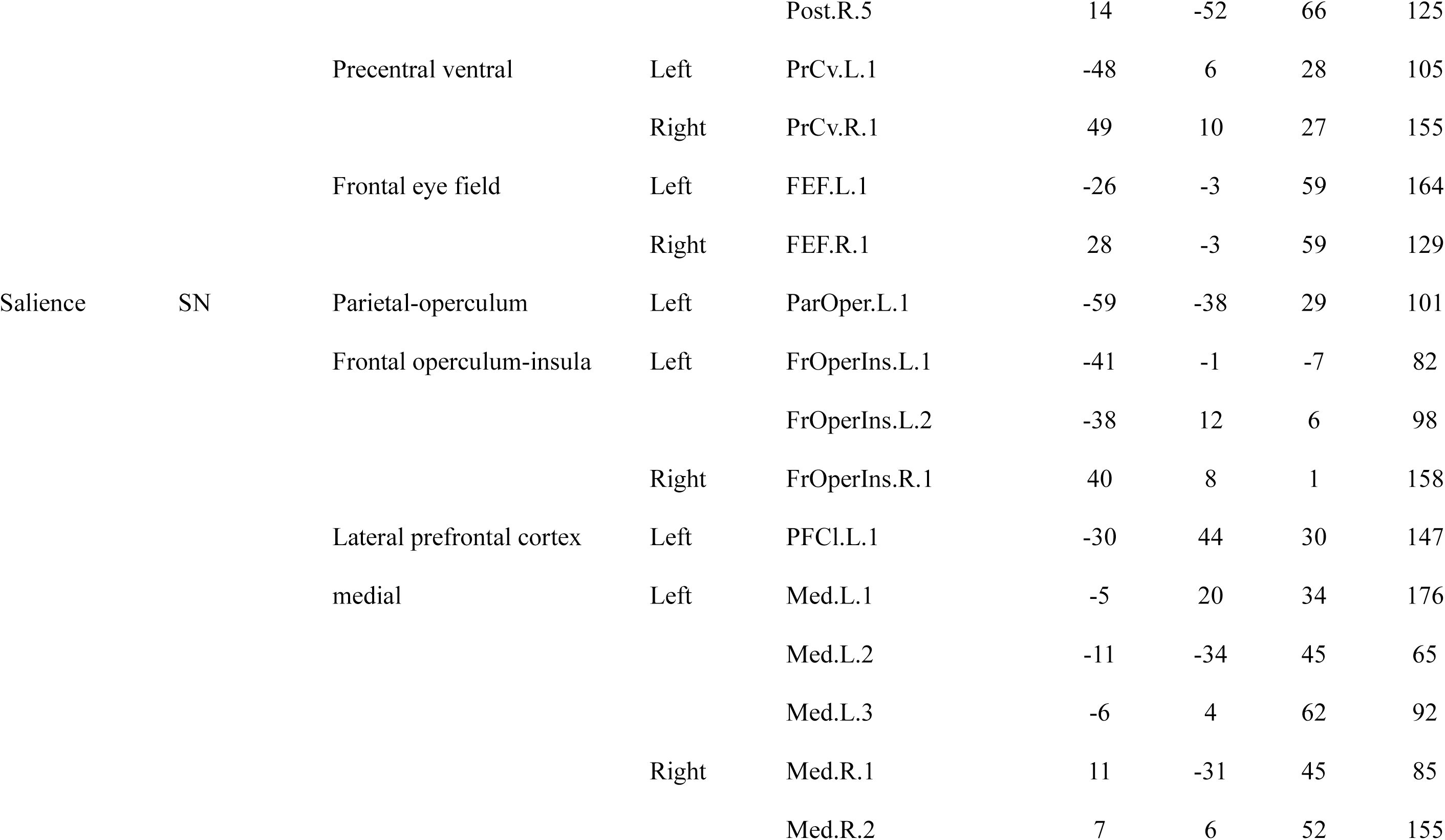
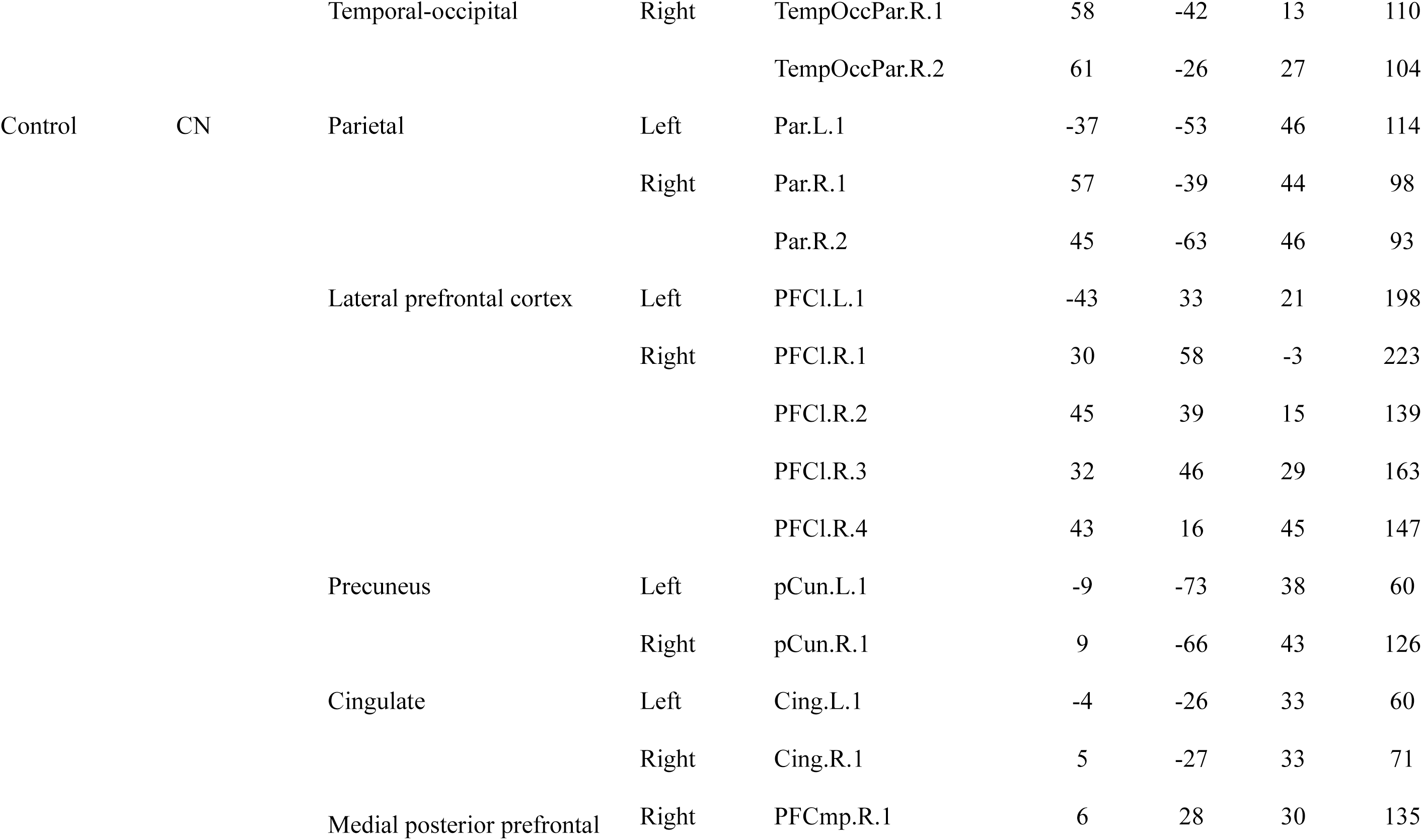
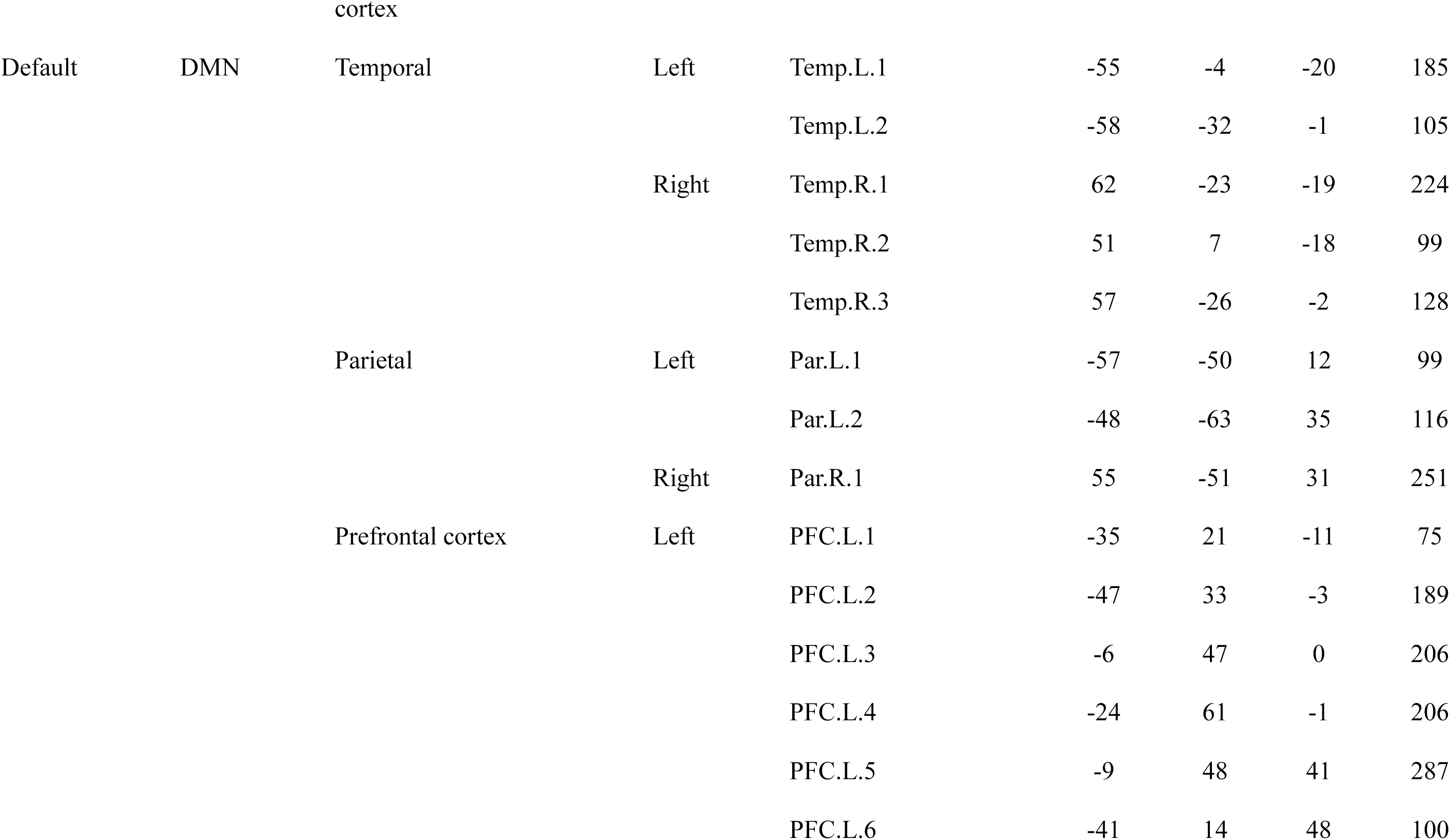
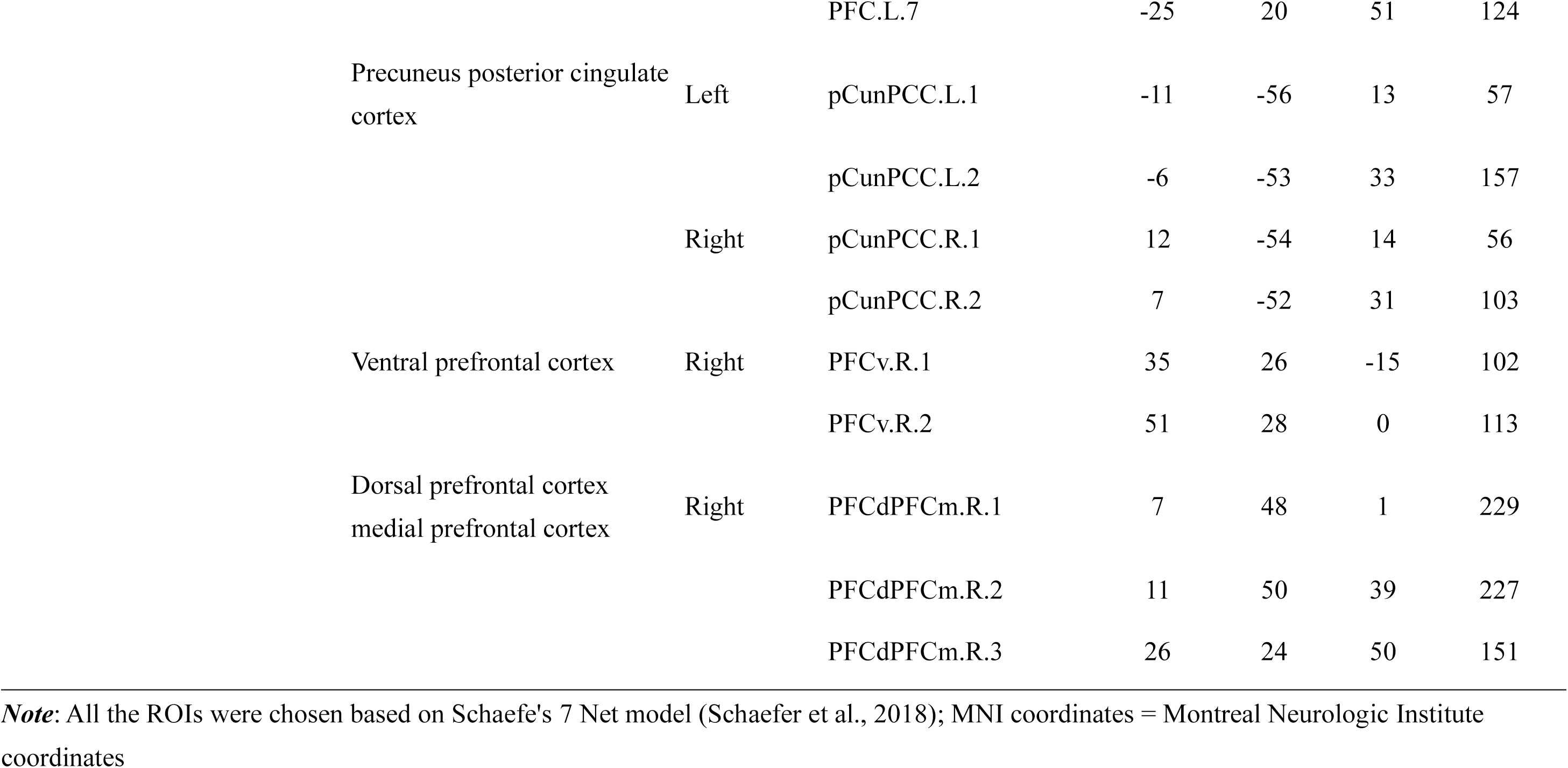
Regions of interest definition and coordinates.

#### 2.6.2 Functional connectivity analyses

The analyzed window corresponded to the period in which the last task image (city or mountain) was presented prior to the thought-probe. This window was defined as the 1000 ms interval immediately before the start of the scrambled image (stimuli image: 100% coherent; scrambled image: 0% coherent) that occurred before the thought probe (Kawashima & Kumano, 2017). Source-estimated connectivity analysis was conducted using this 1000 ms duration. Each participant’s data comprised 27 1000 ms blocks, which were categorized as either the on-task or MW state based on their level of focus during the in-task thought probe.

Functional connectivity was computed between each ROI pair (i.e., edge) using the weighted phase lag index (wPLI; (Vinck et al., 2011)), a phase-based metric designed to capture consistent non-zero-lag relationships while minimizing the impact of volume conduction. Each ROI consisted of multiple source-reconstructed signals from cortical voxels, which were averaged to generate a single representative time course per ROI. To extract frequency-specific phase information, the Hilbert transform was applied to each ROI time series across four bands: delta (2–4 Hz), theta (4.5–7 Hz), alpha (7.5–12 Hz), and beta (13–34 Hz). Pairwise wPLI was calculated over the 1000 ms connectivity analysis window, yielding a symmetrical connectivity matrix for each participant, frequency band, and state.

### 2.7 Statistical analysis

After the connectivity calculation, each participant’s data contained the on-task state, deliberate MW state, and spontaneous MW state wPLI (2016 (combination of 64 ROIs) × 4 frequencies × 3 states = 24192). To identify which ROI pairs exhibited differing wPLI values across the three states, we applied the network-based statistic (NBS) (Zalesky et al., 2010), a nonparametric multiple comparison procedure for testing differences in brain network connectivity, for each frequency band. In line with several previous studies (DeSerisy et al., 2021; Kreilkamp et al., 2021; Lopes et al., 2017; Mehraram et al., 2025; Nelson et al., 2017; Wang et al., 2023; Zhan et al., 2019), we performed NBS analyses across a range of primary thresholds to assess the robustness of the results. The threshold was determined based on stability and conservative considerations, influencing only the sensitivity of the analysis without compromising its accuracy (Zalesky et al., 2010). First, to identify supra-threshold edges, we applied an F-statistic threshold F_primary_ from 5 (df 1= 2, d 2 = 60, p < 0.009) to 7 (df 1= 2, df 2 = 60, p < 0.002) in 0.1 increments. Within this range, values from 5.9 to 6.1 demonstrated stability and conservativeness, leading to the application of F_primary_ at 6.1(df 1= 2, df 2 = 60, p < 0.003) (for details, see Supplementary Material Table 1). Suprathreshold edges were grouped into clusters, defined as connected components of edges sharing at least one common node during the NBS. We performed a permutation test with 20,000 iterations to determine the statistical significance of each cluster. In each permutation, condition labels were shuffled, and the maximum cluster size was recorded to generate an empirical null distribution. Observed cluster sizes were then compared to this distribution to obtain corrected p-values. Finally, component-wise p-values were corrected for multiple comparisons using the Bonferroni method to control the family-wise error rate (FWER).

Given the analysis across four frequency bands, a corrected significance threshold of α = 0.0125 (0.05/4) was applied. This procedure was applied separately to each of the four frequency bands (delta, theta, alpha, beta). We then applied a t-statistic-based NBS to these clusters using threshold T_primary_ from 2 (df = 30, p < 0.03, one-tailed) to 4 (df = 30, p < 0.0001, one-tailed) in 0.1 increments. Six contrasts were tested: [deliberate MW > on-task], [deliberate MW < on-task], [spontaneous MW > on-task], [spontaneous MW < on-task], [deliberate MW > spontaneous MW], and [deliberate MW < spontaneous MW]. Relatively stable edges were observed across thresholds of 2.0–2.5 for [deliberate MW > spontaneous MW] and 2.3–2.6 for [deliberate MW > on-task] within threshold T_primary_ from 2 to 4. Hence, we set the contrast T_primary_ at 2.5 (df = 30, p < 0.008, one-tailed), as it overlapped across the two comparisons and served as a conservative threshold (for details, see Supplementary Material Tables 2 and 3). Component-wise p-values were corrected for multiple comparisons using the Bonferroni method to control the FWER. Given six independent tests, a corrected significance threshold of α = 0.0083 (0.05/6) was applied. After ROI pairs showing significant differences in wPLI values between deliberate MW and spontaneous MW were identified based on NBS, the corresponding wPLI values for each state were extracted, and Spearman rank correlations were performed with attentional trait measures. These included nine attention-related measures, subscales from the CAARS, and the MWS (MWS-D and MWS-S). In total, we conducted correlation analyses across 7 ROI pairs that showed significant differences specifically between deliberate and spontaneous MW, examining their associations with six attentional traits measures. To correct for multiple comparisons, the Benjamini–Hochberg false discovery rate (FDR) correction (Benjamini & Hochberg, 1995) was applied with a significance threshold set at q = 0.05.

## 3. Results

Of the 44 participants initially recruited, three were excluded due to equipment malfunctions (n = 2) or for falling asleep during the experiment (n = 1). An additional three participants provided only “on-task” or “MW” responses on the thought probes, and seven others responded exclusively with either “deliberate MW” or “spontaneous MW,” resulting in missing data for at least one condition. Because the NBS method requires complete data across all conditions, we restricted our analysis to the 31 participants who provided at least one valid report for all three cognitive states: deliberate MW, spontaneous MW, and on-task.

### 3.1 Behavior results

The descriptive statistics for the questionnaire measures are presented in Table 2.

**Table 2.** Descriptive statistics for attentional traits-related questionnaires.

|  | Mean | SD | Kurtosis | Skewness |
| --- | --- | --- | --- | --- |
| **CAARS** | 55.36 | 10.97 | 2.73 | 0.45 |
| **A** |  |  |  |  |
| **CAARS** | 59.16 | 11.51 | 2.25 | 0.30 |
| **B** |  |  |  |  |
| **CAARS** | 52.00 | 9.47 | 2.13 | 0.14 |
| **C** |  |  |  |  |
| **CAARS** | 54.71 | 10.34 | 2.96 | 0.36 |
| **H** |  |  |  |  |
| **MWS-D** | 16.58 | 5.69 | 2.41 | -0.34 |
| **MWS-S** | 16.52 | 5.29 | 2.64 | -0.56 |

**Note:** CAARS A, Conners' Adult ADHD Rating Scale inattention/memory subscale T-score; CAARS B, Conners' Adult ADHD Rating Scale hyperactivity/restlessness subscale T-score; CAARS C, Conners' Adult ADHD Rating Scale impulsivity/emotional lability subscale T-score; CAARS H, Conners' Adult ADHD Rating Scale ADHD Index T-score; MWS-D, Mind-Wandering Scale: Deliberate Mind-Wandering; MWS-S, Mind-Wandering Scale: Spontaneous Mind-Wandering.

The average RT representing processing speed during the gradCPT task was 0.85 s (SD = 0.10). The mean of standard deviations of RTs, an indicator of attentional stability and response variability, was 0.17 (SD = 0.06). Participants exhibited a low average error rate of 0.04 (SD = 0.04).

During the task, participants reported an average of 3.97 episodes of deliberate MW (SD = 3.82) and 7.87 episodes of spontaneous MW (SD = 4.90) out of 27 in-task thought probes, while the remaining 15.16 probes (SD = 5.68) were classified as on-task (see Figure 3).

**Figure 1.**
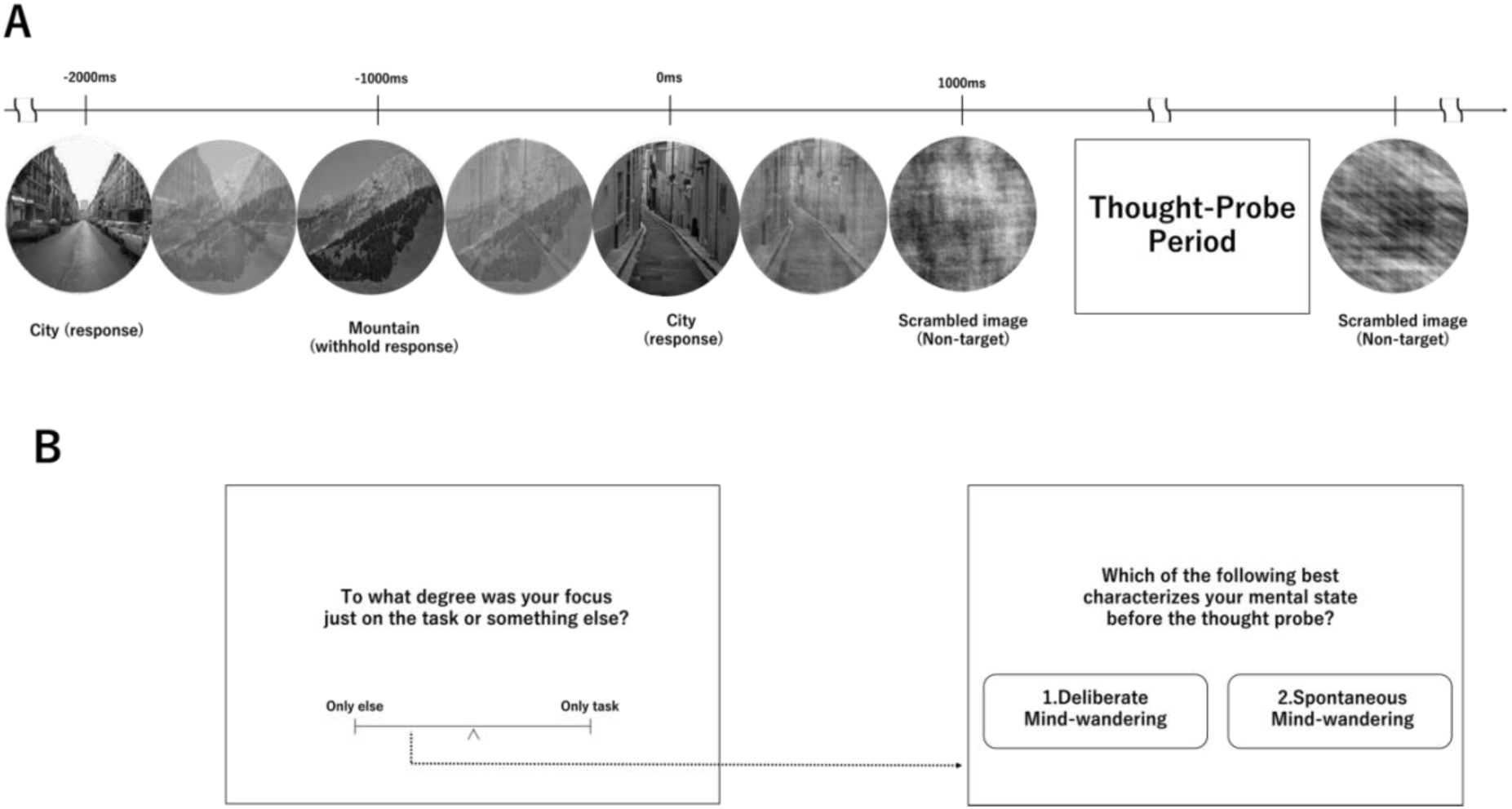
GradCPT paradigm and classification of cognitive states from thought probes. (A) Gradual-onset continuous performance task (gradCPT) paradigm. Participants viewed gradually changing images of city (frequent) and mountain (rare) scenes and were instructed to respond to city images but withhold responses to mountain images. A thought probe appeared every 30–60 s. (B) Classification of cognitive states based on thought probe responses. Using a continuous scale (0 = only task, 100 = only else), participants rated the extent to which their thoughts were focused on the task. If their rating indicated off-task focus (≤ 50), their state was further classified as deliberate or spontaneous MW, whereas on-task ratings (> 50) were classified as the on-task state.

**Figure 2.**
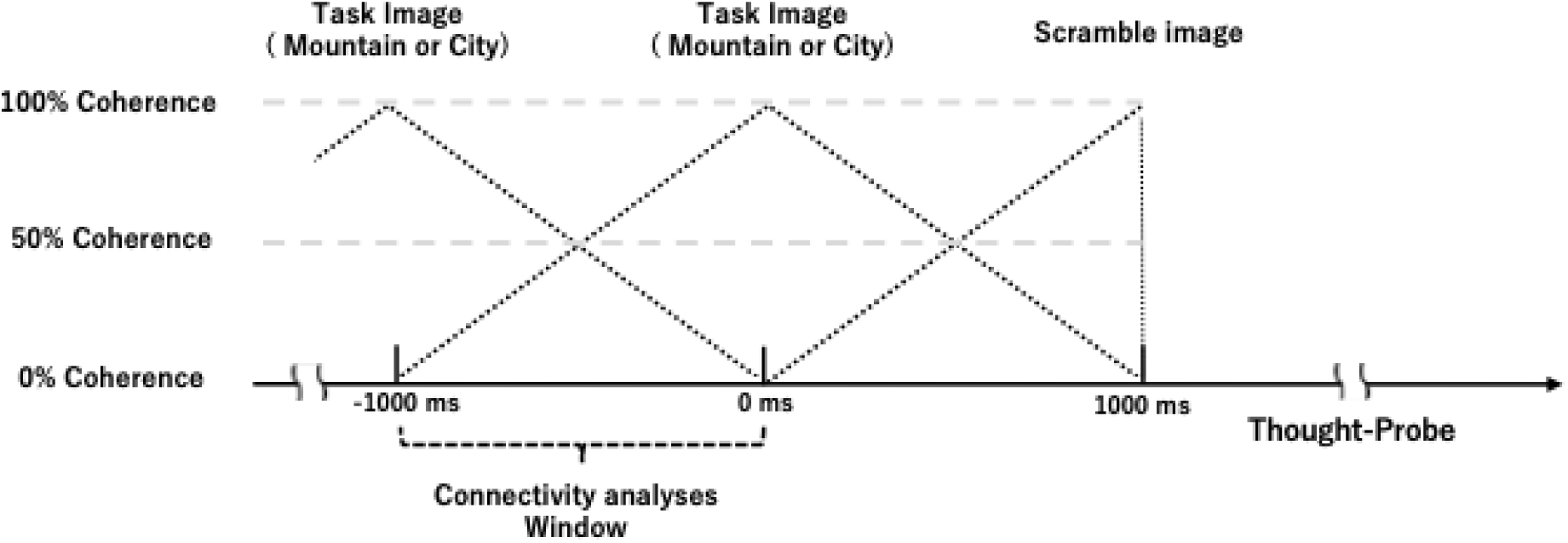
Connectivity analysis window. The 1000 ms period preceding the onset of the scrambled image (-1000 ms to 0 ms from the onset of the scrambled image), during which only the task image (either city or mountain) was presented, was used for source-estimated connectivity analysis.

**Figure 3.**
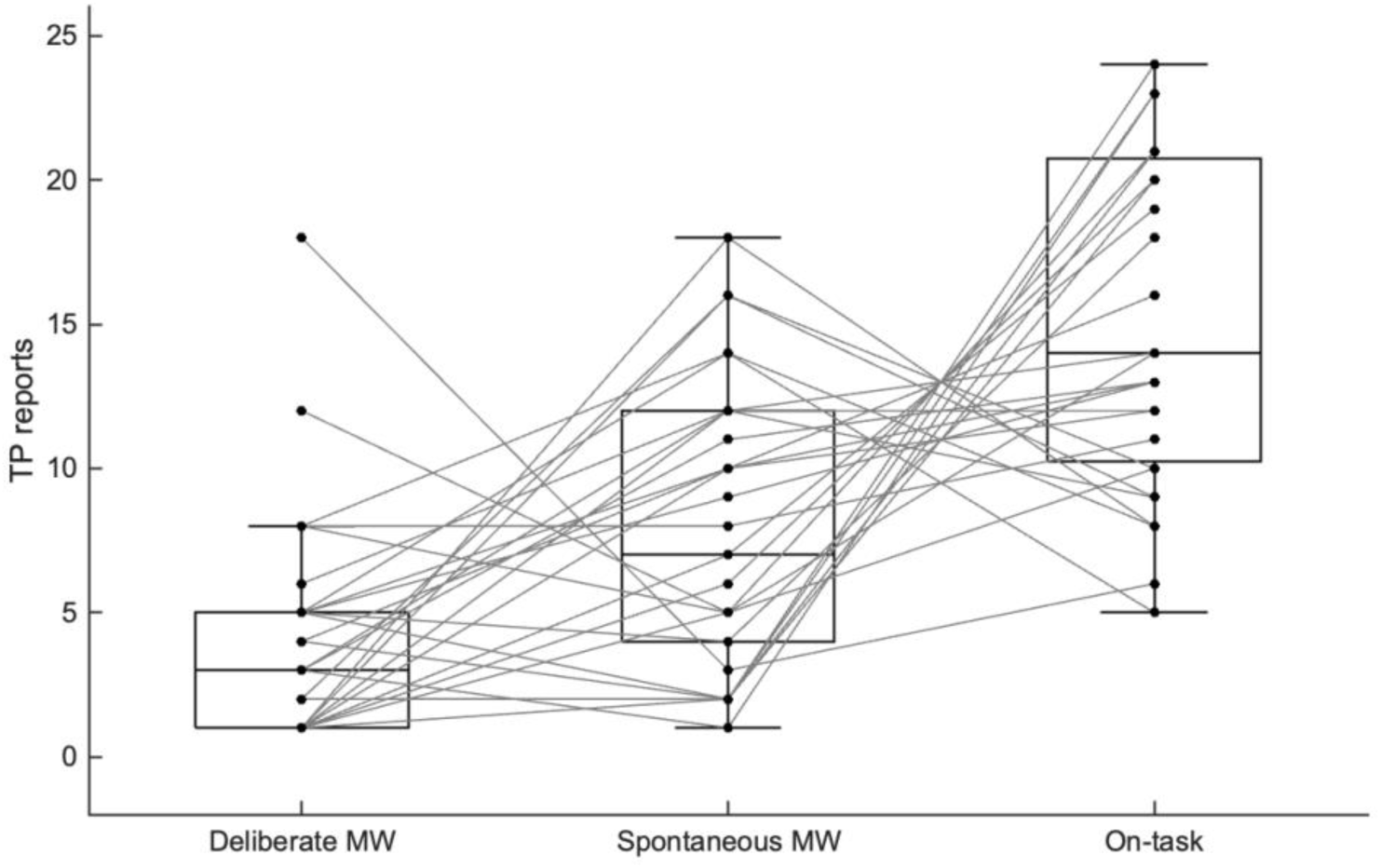
Number of deliberate and spontaneous mind-wandering and on-task states identified from in-task thought probes during the gradCPT. (total = 27 probes).

### 3.2. FC differences between the MW and on-task states

#### 3.2.1 Three-state F-statistic-based NBS result

Based on F-statistic-based NBS, connectivity clusters showing a significant main effect (NBS-adjusted p < .0125, Bonferroni-corrected for four frequencies) were extracted, indicating that the null hypothesis of equal connectivity across all three conditions was rejected. Significant wPLI edges were identified at the alpha band (7.5–12 Hz) (10 edges, 11 ROIs) (for details, see Supplementary Material Figure S9.).

#### 3.2.2 T-statistic-based NBS result

Based on the significant clusters identified through the F-statistic-based NBS, we extracted the corresponding wPLI values and conducted t-statistic-based NBS. These follow-up t-statistic-based NBS analyses identified the specific contrasts that accounted for the observed main effects.

##### 3.2.2.1 T-statistic-based NBS result between deliberate MW and spontaneous MW

We conducted t-statistic-based NBS analyses at threshold of t = 2.5 (df = 30, p < 0.01, one-tail), comparing deliberate and spontaneous MW. Deliberate MW showed significantly stronger wPLI connectivity than spontaneous MW at the alpha band (NBS-adjusted *p* < 0.0083, Bonferroni-corrected for six contrasts).

In the alpha band, one significant cluster was observed (Figure 4), comprising seven edges and showing increased connectivity between the DMN (two ROIs), CN (four ROIs), and SN (two ROIs). This cluster was centered on the right frontal operculum–insula of the SN, which showed increased FC with the left medial region of SN, left precuneus posterior cingulate cortex of the DMN, left and right precuneus region of the CN, and right cingulate region of the CN, which also exhibited greater FC with the DMN left precuneus posterior cingulate cortex region.

**Figure 4.**
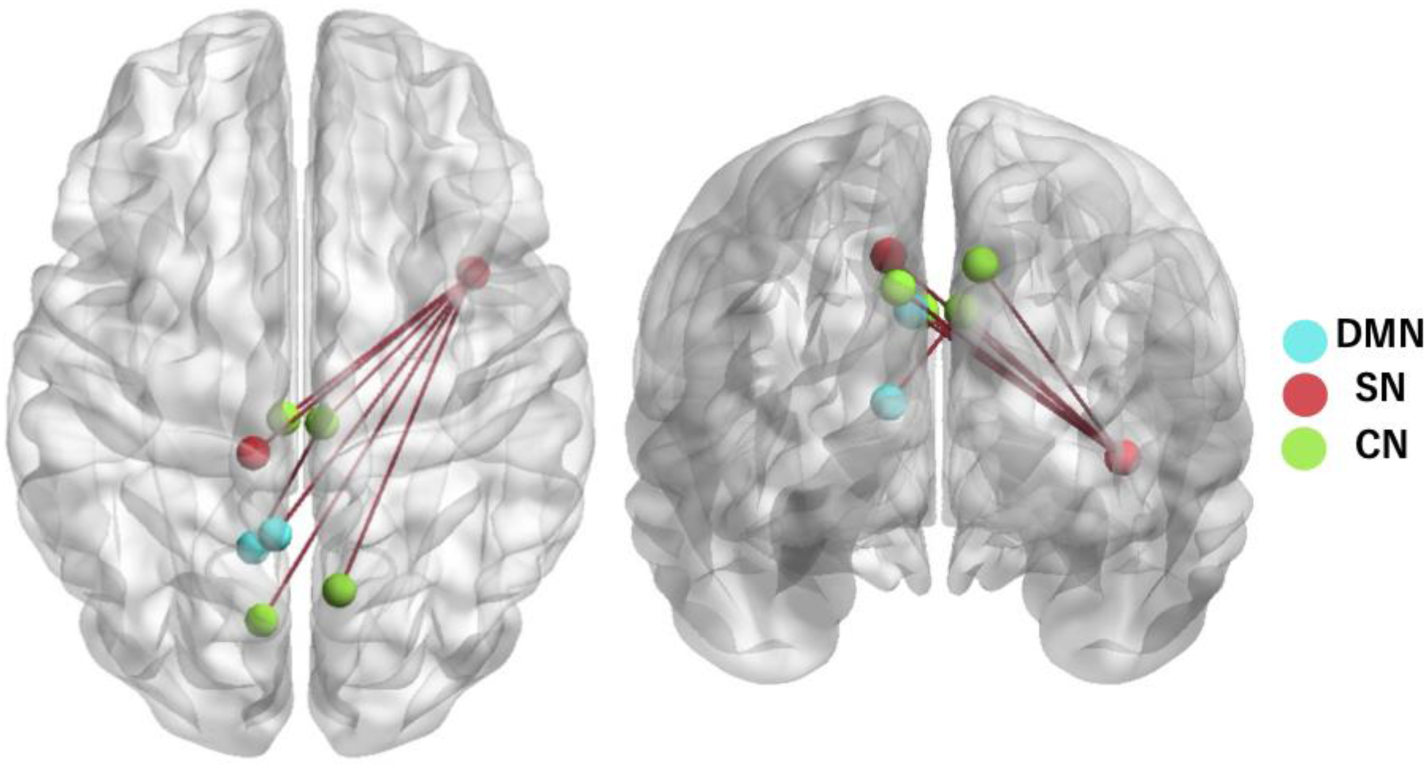
Alpha band T-statistic-based NBS result between deliberate MW and spontaneous MW. Relative to spontaneous MW, the deliberate MW state shows one significant greater wPLI cluster in the alpha band, comprising seven edges. This cluster included SN right frontal operculum–insula centered connections between the DMN (two ROIs), CN (four ROIs), and SN (two ROIs). DMN, default mode network; SN, salience network; CN, control network

No significant connectivity differences were observed between deliberate and spontaneous MW in the delta, theta, or beta bands. Meanwhile, no connectivity clusters showed significantly greater connectivity during spontaneous MW compared with deliberate MW at the same primary threshold.

##### 3.2.2.2 T-statistic-based NBS result between deliberate MW and on-task state

We investigated differences in wPLI between deliberate MW and the on-task state at NBS threshold t = 2.5 (df = 30, p < 0.01, one-tail). Deliberate MW showed significantly stronger connectivity compared with the on-task state at the alpha band (NBS-adjusted p < 0.0083, Bonferroni-corrected for six contrasts). The alpha band showed elevated connectivity across seven edges, including the DMN (two ROIs), CN (three ROIs), SN (two ROIs), and DAN (one ROI). In this cluster, the right frontal operculum–insula of the SN was the center node, which showed greater FC to the left medial region of the SN, right cingulate region of the CN, left and right precuneus region of the CN, left precuneus posterior cingulate cortex of the DMN, and left posterior region of the DAN, which also showed greater FC with the DMN left parietal region (see Figure 5).

**Figure 5.**
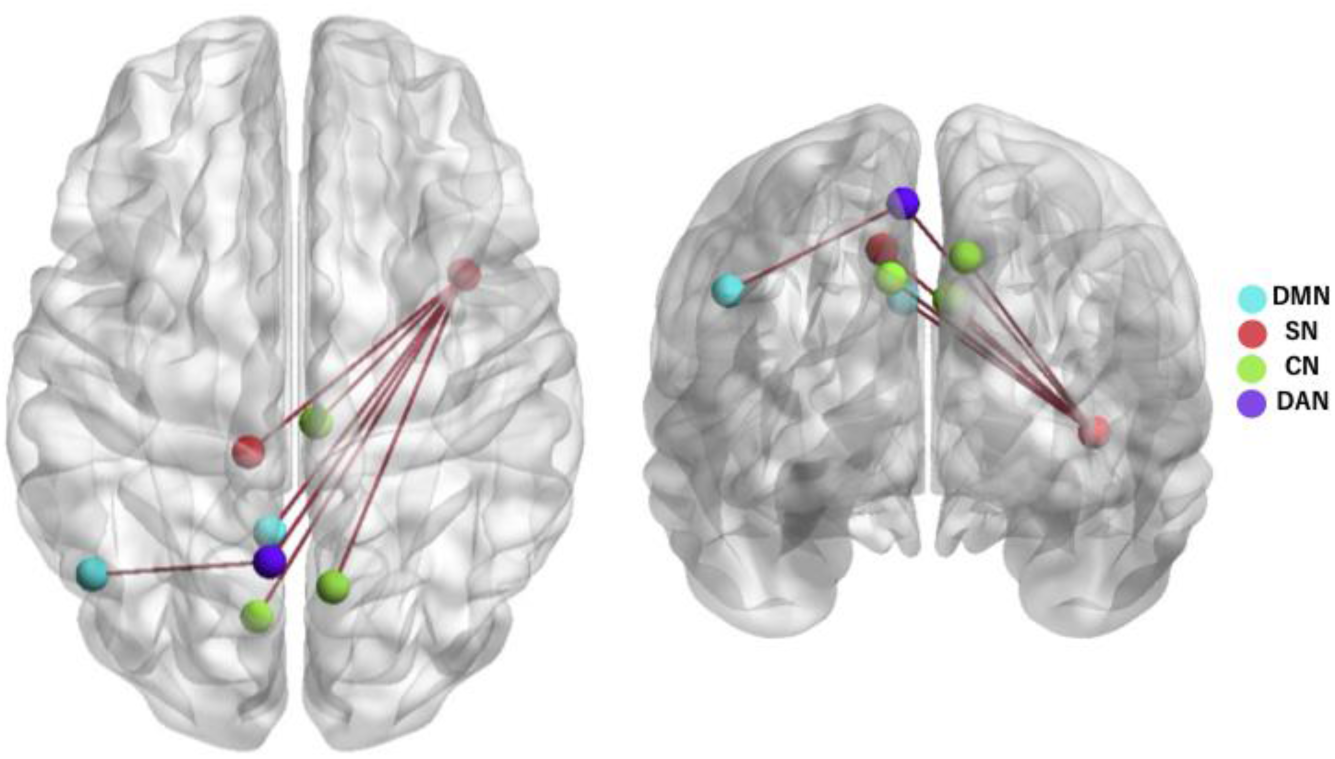
T-statistic-based NBS result between deliberate MW and on-task state at the alpha band (7.5–12 Hz). The cluster included SN right frontal operculum–insula centered connections. This cluster contained seven greater wPLI edges across the DMN (two ROIs), CN (three ROIs), SN (two ROIs), and DAN (one ROI) during deliberate MW compared with the on-task state. DMN, default mode network; SN, salience network; CN, control network; DAN, dorsal attention network.

No significant connectivity differences were observed between deliberate and on-task state in the delta, theta, or beta bands. No connectivity clusters showed significantly higher wPLI during the on-task state compared with deliberate MW at the same primary threshold.

##### 3.2.2.3 T-statistic-based NBS result between spontaneous MW and the on-task state

No significant connectivity differences were found between spontaneous MW and the on-task state in either contrast at the threshold of t = 2.5 (df = 30, p < 0.01, one-tail).

### 3.3 Relationships between FC and attentional tendencies

Correlation analyses indicated no statistically significant associations between functional connectivity during either deliberate MW or spontaneous MW with any of the attention-related measures examined, following the application of FDR correction.

## 4. Discussion

We investigated how large-scale network connectivity differs during deliberate MW, spontaneous MW, and on-task states using EEG source-level functional connectivity alongside on-line experience sampling during a sustained attention task. By dissociating deliberate and spontaneous MW at the state level, our findings reveal a clear neural distinction between these two forms of internally directed cognition. Our results reveal that (1) compared with spontaneous MW, deliberate MW showed an alpha-band connectivity cluster centered on the right frontal operculum–insula of the SN, linking regions of the SN, CN, and DMN, while the connected right cingulate region of the CN exhibited greater FC with the DMN left precuneus posterior cingulate cortex region. (2) Compared with the on-task state, deliberate MW showed an alpha-band connectivity cluster centered on the right frontal operculum–insula of the SN, linking regions of the SN, CN, and DMN, while the connected left posterior DAN exhibited greater FC with the left parietal region of the DMN.

Our findings suggest that relative to spontaneous MW and the on-task state, deliberate MW is characterized by hub-like involvement of the right frontal operculum−insula within the SN. The SN has been linked to the process of detecting salient sensory inputs or cognitively relevant internal events (Uddin, 2015). In particular, the anterior insula of the SN has been described as a switching hub that reallocates processing resources between externally oriented and internally directed cognition (Blomberg et al., 2022; Goulden et al., 2014; Li et al., 2021; Menon & Uddin, 2010; Seeley, 2019; Uddin, 2015). Increased SN−DMN coupling has been associated with a higher MW tendency (Blomberg et al., 2022; Webb et al., 2021). By contrast, focused task engagement is typically characterized by positive SN−CN coupling (Denkova et al., 2019; Goulden et al., 2014; Menon & Uddin, 2010). However, these patterns do not fully capture cognitive states in which internal cognition is intentionally maintained. Our results show that deliberate MW is characterized by coordinated connectivity between the SN FrOperIns and both the CN and DMN. Notably, the cingulate region of the CN that was coupled with the SN FrOperIns also showed increased connectivity with the posterior cingulate cortex of the DMN relative to spontaneous MW. Previous work has linked coordinated DMN−CN coupling with higher deliberate MW traits (Golchert et al., 2017), meditation with intentional attentional control (Marzetti et al., 2014), and future-oriented planning (Maillet et al., 2019), suggesting that intentionally oriented internal thought may involve coordinated interactions between these networks. Additionally, fMRI evidence indicates that SN–DMN activity is anticorrelated during sustained attention and boredom, whereas it exhibits correlated activity during interest induction(Danckert & Merrifield, 2018). The observation of this coupling, together with enhanced SN−CN and SN−DMN coupling, suggests that deliberate MW recruits control processes supporting the goal-directed regulation of internal thought through SN-centered integration of control and default mode systems. Accordingly, our results extend SN, DMN, and CN interaction models by demonstrating an integrative role of the SN in intentional MW.

Interestingly, relative to the on-task state, deliberate MW additionally showed enhanced SN–DAN–DMN coupling. DAN is involved in top-down attentional control (M. D. Fox et al., 2006), supporting the prioritization of attentional resources (Szczepanski et al., 2013) and transient shifts toward off-task thought (Turnbull et al., 2019). DAN and DMN often exhibit opposing dynamics, with externally oriented tasks engaging the DAN and internally directed thought engaging the DMN (Kucyi et al., 2020, 2021; Maillet et al., 2019; Mittner et al., 2014). However, increased coupling between the DMN, DAN, and SN has been observed during meditation, a state characterized by the intentional regulation of attention and control of thought flow (Froeliger et al., 2012). The left posterior DAN has been linked to top-down attentional control (Lee et al., 2023; Machner et al., 2022; Rohr et al., 2017; Szczepanski et al., 2013; Wen et al., 2012), while the left parietal DMN supports introspective and memory-related processes (Menon, 2023; Sestieri et al., 2011; Smith et al., 2021). Collectively, given that the right frontal operculum–insula of the SN might serve as a hub integrating internally and externally oriented processes, the observed SN−DAN−DMN coupling during deliberate MW may support the intentional regulation of attentional resources between internal and external systems.

Moreover, our findings provide functional evidence that deliberate MW is specifically supported by alpha band network interactions. Alpha band activity has long been associated with selective attention (Sauseng et al., 2005), particularly the suppression of task-irrelevant information (Foxe & Snyder, 2011; Kam et al., 2022; Klimesch et al., 2007).

Previous research has proposed that alpha band inter-regional coupling might support internal information processes in working memory (Palva & Palva, 2011) and meditation with controlled attention resources (Marzetti et al., 2014). Although these roles may appear contradictory, they can be reconciled by considering the intentional nature of deliberate MW. Specifically, alpha band coupling may simultaneously inhibit externally oriented processing while promoting coordinated interactions among networks that support internally generated thought. Consistent with this view, changes in alpha band amplitude have been repeatedly identified as neural markers of MW (Braboszcz & Delorme, 2011; Compton et al., 2019; Rodriguez-Larios & Alaerts, 2021; van Son, De Blasio, et al., 2019; van Son, de Rover, et al., 2019), and greater alpha power or alpha band phase locking has been reported during deliberate MW (Kam et al., 2021; Martel et al., 2019). Taken together, our findings indicate that the oscillation of alpha band inter-network connectivity observed during deliberate MW may reflect the intentional regulation of internal cognition.

In this study, no significant FC differences were observed between spontaneous MW and the on-task state during the sustained attention task. Within the sensitivity of the current EEG source-level connectivity approach, this finding suggests that spontaneous MW may not be reliably distinguished from on-task focus based on network connectivity. Prior studies have reported that left-lateralized structural variability in default mode regions is observed with higher spontaneous MW tendencies (Golchert et al., 2017). Beyond structure, temporal variability of the resting state DMN activity has also been found to be correlated with spontaneous MW traits (Sorella et al., 2025). One possible interpretation, informed by prior trait-level and resting-state studies, is that spontaneous MW may be more strongly reflected in structural characteristics or temporal variability within DMN subregions. Importantly, this possibility cannot be directly tested with the present data, which focused on phase-based connectivity measures. To clarify these aspects, future work should employ MRI to assess activity differences within specific DMN subregions, enabling a clearer characterization of how spontaneous MW diverges from task-focused states.

There are several limitations that should be considered when interpreting the results of the current study. One important limitation concerns the spatial precision of EEG source localization, particularly for deep cortical regions such as insula and precuneus regions. EEG source estimation inherently has lower spatial resolution than hemodynamic imaging methods, and its accuracy can be influenced by preprocessing procedures, head model assumptions, and source estimation methods (Asadzadeh et al., 2020; Gomez-Tapia et al., 2025; Michel & Brunet, 2019). To mitigate these issues, several methodological precautions were implemented. We utilized a 64-channel EEG system, which offers relatively high accuracy in source estimation (Shen et al., 2023; Sohrabpour et al., 2015). Source reconstruction was performed using a boundary element method (BEM) head model implemented in OpenMEEG, which provides a more anatomically realistic approximation of head conductivity than simplified spherical models, despite not incorporating individual MRI data (Chowdhury et al., 2015; Gramfort et al., 2010; Qin et al., 2023; Stropahl et al., 2018). To further account for interindividual variability in cortical geometry (Ahlfors et al., 2010), unconstrained dipole orientations were used and flattened using PCA. Additionally, functional connectivity was quantified using the wPLI to minimize spurious coupling due to volume conduction (Vinck et al., 2011). Importantly, the observed alpha band inter-network coupling involving the precuneus region and frontal operculum insula is consistent with prior EEG, magnetoencephalography (MEG), and fMRI findings linking these regions to self-referential processing (Bao & Frewen, 2022), controlled attention during meditation (Marzetti et al., 2014), and MW (Kucyi et al., 2016, 2024). This convergence supports the plausibility of our network-level interpretation of deliberate MW despite the spatial limitations inherent to EEG source estimation. Another limitation concerns the NBS approach, which requires the selection of a primary threshold and is inherently arbitrary. However, this arbitrariness affects only the sensitivity of the analysis and does not compromise its accuracy (Zalesky et al., 2010). To ensure rigor, we adopted stepwise sensitivity analysis using multiple primary thresholds and selected primary thresholds based on stability and conservative considerations, as commonly employed in previous studies (DeSerisy et al., 2021; Mason et al., 2023; Nelson et al., 2017; Pua et al., 2018; Wagner et al., 2019; Wang et al., 2023; Yang et al., 2018). This conservative choice yielded strong, topologically focal differences, supporting the reliability and validity of our findings despite methodological constraints. Nonetheless, the direction and causality of these network interactions remain unclear. Future work using causal approaches such as cognitive training or frequency-specific neuromodulation (e.g., tACS) may help elucidate these mechanisms.

In conclusion, this study provides state-level neural evidence for intentional forms of MW. Using source-level EEG connectivity and in-task experience sampling, we found that deliberate MW was characterized by alpha band coupling centered on the right frontal operculum–insula of the SN, linking the SN with executive control and default mode systems relative to both spontaneous MW and on-task states. Additionally, increased connectivity between the SN, DAN, and DMN was observed during deliberate MW compared with the on-task state. These findings extend prior trait-level research by revealing that intentionality exhibits distinct state-level network interactions, emphasizing the neural signature of internally directed cognition. More broadly, our results support the view that deliberate MW reflects a controlled mode of internal thought, emerging from the integration of self-referential processing with executive and attentional regulation.

## Data and code availability

All EEG and gradCPT data generated or analyzed in this study are accessible via the Open Science Framework (OSF) below. https://osf.io/37u9h/overview?view_only=fef57a2dd6ba4257bba85bd127b24d32

## Author contributions

X.Y. and R.O. designed the experiments and prepared the manuscript. X.Y. and M.I. wrote the gradCPT code. X.Y. collected and analyzed the data. Y.K., T.T., and R.O. provided important suggestions on data analysis and the manuscript. X.Y., T.T., and R.O. wrote the paper.

## Acknowledgments

We express our gratitude to Michael Esterman and Aaron Kucyi for providing the original code for the gradual continuous performance tasks, which served as a reference for our implementation. We also extend our thanks to all the participants involved in the study.

