## Supplementary Materials for "Large-scale brain network differences during deliberate and spontaneous mind-wandering in a sustained attention task: An electroencephalography source-level connectivity study"

***Task instruction***

Text in *italics* indicates the instructions as they were given to participants.
All task instructions and example screens were presented and explained in Japanese.

### Step 1. Task stimulus presentation

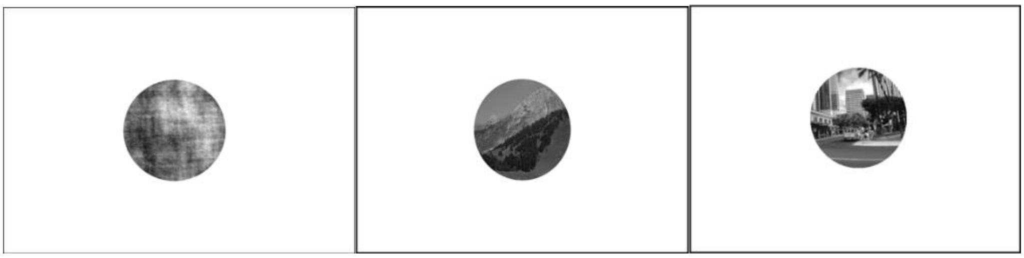

**Figure S1. Example of task stimuli shown to participants**. All images used in the experiment (city, mountain, and scrambled images) were shown once to each participant. All example images were displayed without gradual transitions (i.e., no fade-in/fade-out).

*During this task, images of mountains and images of cities will be presented on the screen. These images will now be shown to you, so please take a moment to familiarize yourself with both the mountain and city images.*

### Step 2. Explanation of the task structures
### Step 2.1 Explanation of task response

*Throughout the task, the mountain and city images will appear alternately. The images will gradually change as they are presented.*

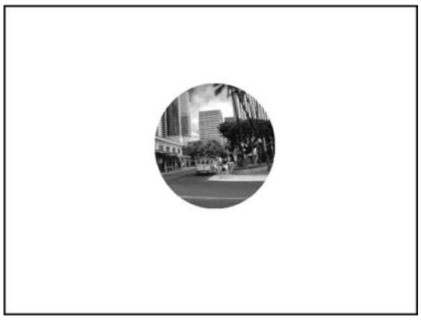

**Figure S2. Example of a city scene stimulus (go trial)**

*When you see a city image, press the space key once with your right hand.*

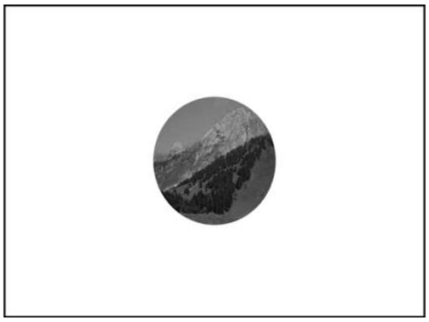

**Figure S3. Example of a mountain scene stimulus (no-go trial)**

*When you see a mountain image, do not press the space key. In most cases, city images will be presented.*

### Step 2.2 Thought-probe during the task

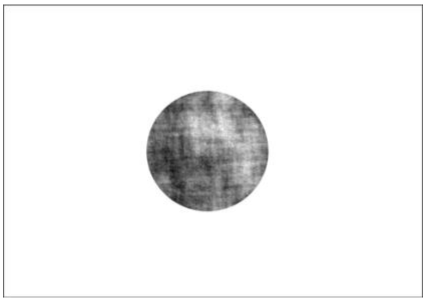

**Figure S4. Example of a scrambled image stimulus**

*At random intervals during the task, a neutral image unrelated to the task (a scrambled image) will be presented. Immediately after this image, a series of questions (thought probe window) will appear. Please answer these questions honestly. Each question will be displayed for 20 s, so be sure to respond within that time.*

#### Step 2.2.1 Thought-probe question 1

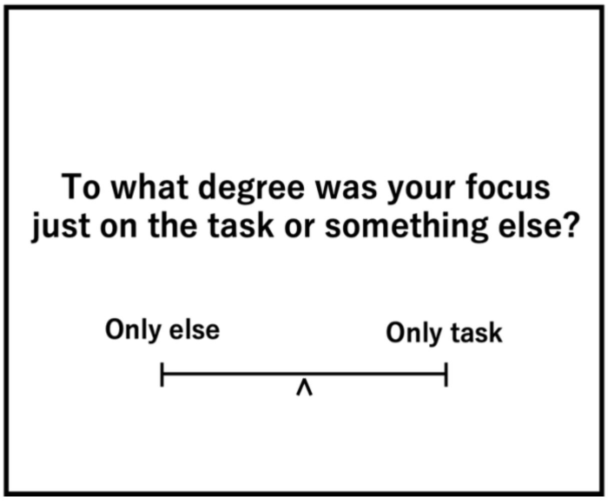


**Figure S5. Example screen for thought-probe question 1**

*The first question asks where your attention was directed immediately before the thought probe period.*
*Use your left hand to select a point on the scale using the "A" (left) and "D" (right) keys to indicate whether your attention was focused on the task or on something else. Press the space key to enter your answer.*

*"Only task" refers to a state in which, immediately before the thought probe period, you were focused on performing the task and were not thinking about anything unrelated to it.*

*"Only task" includes various task-related thoughts, such as:*
*• thoughts about how well you are performing (task performance),*
*• thoughts about the images being presented,*
*• thoughts related to your key presses, etc.*
*Even if you had such thoughts, this still counts as "only task" although the level may vary.*

*"Only else" indicates mind-wandering, that is, thinking about content entirely unrelated to the task immediately before the question period.*
*Examples of mind-wandering include thoughts about what you will eat for dinner, plans with friends, or an upcoming exam.*

### Step 2.2.2 Thought-probe question 2

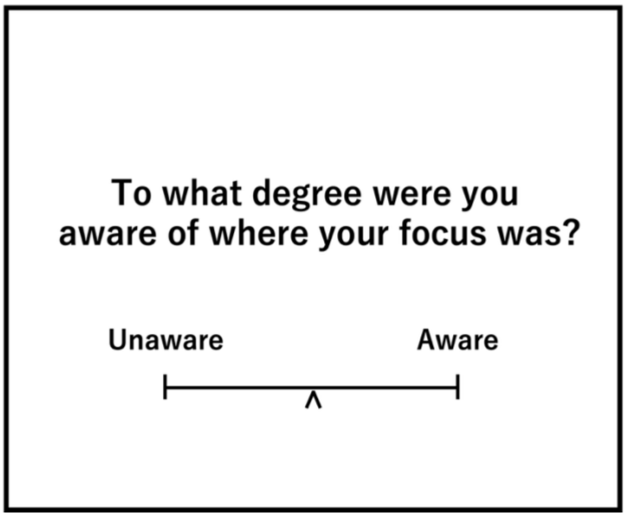


**Figure S6. Example screen for thought-probe question 2**

*The second question asks to what extent you were aware of where your attention was directed immediately before the thought probe period.*

*Use the "A" and "D" keys to select a point on the scale indicating whether you were "aware" or "unaware" and press the space key to enter your answer.*

*For example, while performing the task, sometimes you may notice that your attention has drifted, while at other times you may not realize this until you are asked.*

### Step 2.2.3 Thought-probe question 3

*The third question is presented only if you answered "only else" in Question 1.*

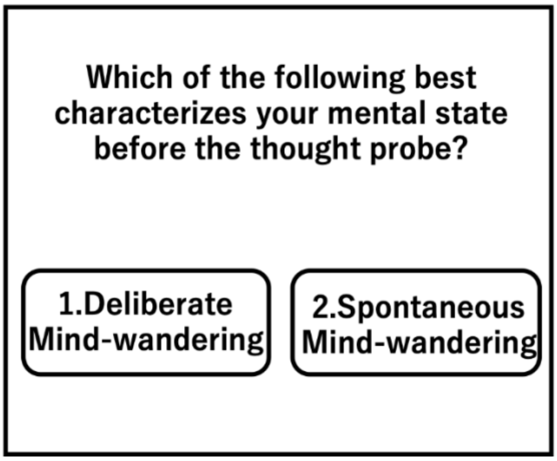


**Figure S7. Example screen for thought-probe question 3**

*Select the option that best describes your mental state immediately before the thought probe period. Respond using the number keys "1" or "2."*

1. *Deliberate mind-wandering*
2. *Spontaneous mind-wandering*

*Deliberate mind-wandering refers to intentionally thinking about things unrelated to the task. Spontaneous mind-wandering refers to unintentionally thinking about task-unrelated matters even though you intended to stay focused.*

*If this question appears, please choose the type of mind-wandering that applies to you.*

### Step 2.2.4 Thought-probe question 4

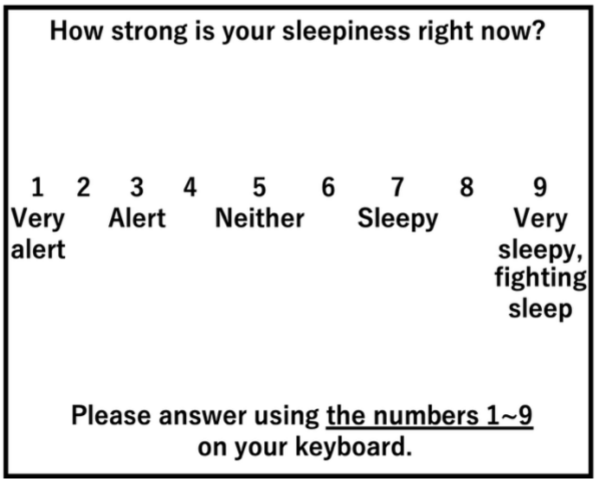


**Figure S8. Example screen for thought-probe question 4**

*The fourth question asks about your level of sleepiness immediately before the thought probe period. Respond using the number keys "1–9" with your left hand.*

*Some response options (2, 4, 6, 8) do not have verbal labels. These represent intermediate levels between the described categories.*
*For example, option "4" corresponds to "somewhat alert."*

### Step 2.3 Practice session 1

*We will now begin the first practice session. The images will gradually change as they are presented.*

*When you see a city image, press the space key once with your right hand.*
*When you see a mountain image, do not press the space key.*
*If a question appears, please respond using your left hand.*
*For practice purposes, when Question 1 appears, please select "only else."*

*During the practice session, task execution and thought probes were explained as needed.

### Step 2.4 Practice session 2

*In this second practice session, the images will be presented at a faster rate.*
*When you see a city image, press the space key once with your right hand.*
*When you see a mountain image, do not press the space key.*
*The task will be more difficult, but please do your best.*
*For practice purposes, when Question 1 appears, please select "only task."*

*During the practice session, task execution and thought probes were explained as needed.

### Step 3 Start of the main experiment

*In the main experiment, the task will proceed at the same speed as in the previous practice.*
*Please do your best. If you make mistakes or miss some images, continue the task without stopping.*
*When a question appears during the task, be sure to answer honestly.*

***F-statistic NBS results for deliberate MW, spontaneous MW, and on-task states***

**Table S1. Alpha band (7.5–12Hz) three-state NBS results (F-statistic) across $F_{primary}$ thresholds from 5 to 6.3 (0.1 increments) (NBS-adjusted p < .0125)**

| **Edges** | | **$F_{primary}$ Threshold** | | | | | | | | | | | | | |
|---|---|---|---|---|---|---|---|---|---|---|---|---|---|---|---|
| ROI 1 | ROI 2 | 5 | 5.1 | 5.2 | 5.3 | 5.4 | 5.5 | 5.6 | 5.7 | 5.8 | 5.9 | 6 | 6.1 | 6.2 | 6.3 |
| SN.ParOper.1.L | CN.Cing.1.L | ○ | ○ | ○ | ○ | ○ | ○ | ○ | ○ | ○ | ○ | ○ | ○ | | |
| SN.ParOper.1.L | SN.Med.2.R | ○ | | | | | | | | | | | | | |
| SN.Med.1.L | DMN.PFC.2.L | ○ | ○ | ○ | ○ | | | | | | | | | | |
| SN.FrOperIns.1.R | CN.Cing.1.L | ○ | ○ | ○ | ○ | ○ | ○ | ○ | ○ | ○ | ○ | ○ | ○ | | |
| SN.FrOperIns.1.R | CN.Cing.1.R | ○ | ○ | ○ | ○ | ○ | ○ | ○ | ○ | ○ | ○ | ○ | ○ | ○ | |
| SN.FrOperIns.1.R | CN.pCun.1.L | ○ | ○ | ○ | ○ | ○ | ○ | ○ | ○ | ○ | ○ | ○ | ○ | ○ | |

| | | | | | | | | | | | | | | |
|---|---|---|---|---|---|---|---|---|---|---|---|---|---|---|
| SN.FrOperIns.1.R | CN.pCun.1.R | ○ | ○ | ○ | ○ | ○ | ○ | ○ | ○ | ○ | ○ | ○ | ○ | ○ |
| SN.FrOperIns.1.R | DMN.Par.1.R | ○ | ○ | ○ | ○ | ○ | ○ | | | | | | | |
| SN.FrOperIns.1.R | DMN.pCunPCC.2.L | ○ | ○ | ○ | ○ | ○ | ○ | ○ | ○ | ○ | ○ | ○ | ○ | ○ |
| SN.FrOperIns.1.R | DAN.Post.5.L | ○ | ○ | ○ | ○ | ○ | ○ | ○ | ○ | ○ | ○ | ○ | ○ | ○ |
| SN.FrOperIns.1.R | SN.Med.1.L | ○ | ○ | ○ | ○ | ○ | ○ | ○ | ○ | ○ | | | | |
| SN.FrOperIns.1.R | SN.Med.2.L | ○ | ○ | ○ | ○ | ○ | ○ | ○ | ○ | ○ | ○ | ○ | ○ | ○ |
| DMN.Temp.2.R | DMN.PFC.7.L | ○ | | | | | | | | | | | | |
| DMN.Temp.1.L | DMN.Par.1.L | ○ | ○ | ○ | | | | | | | | | | |
| DMN.PFC.6.L | CN.pCun.1.L | ○ | ○ | ○ | ○ | | | | | | | | | |

| | | | | | | | | | | | | | | |
|---|---|---|---|---|---|---|---|---|---|---|---|---|---|---|
| DMN.pCunPCC.2.R | CN.Cing.1.R | ○ | ○ | ○ | ○ | | | | | | | | | |
| DMN.pCunPCC.1.L | CN.Cing.1.R | ○ | ○ | ○ | ○ | ○ | ○ | ○ | ○ | ○ | ○ | ○ | ○ | ○ |
| DMN.Par.1.R | CN.PFCl.1.R | ○ | ○ | ○ | ○ | ○ | ○ | | | | | | | |
| DMN.Par.1.R | DMN.PFCv.1.R | ○ | ○ | ○ | ○ | ○ | ○ | | | | | | | |
| DAN.Post.5.L | DMN.Par.2.L | ○ | ○ | ○ | ○ | ○ | ○ | ○ | ○ | ○ | ○ | ○ | ○ | ○ |
| DAN.Post.5.L | DMN.Temp.2.R | ○ | ○ | ○ | ○ | | | | | | | | | |
| DAN.Post.1.L | DMN.Par.1.L | ○ | ○ | ○ | | | | | | | | | | |
| DAN.Post.1.L | DAN.FEF.1.L | ○ | ○ | ○ | | | | | | | | | | |
| DAN.FEF.1.L | DMN.Par.1.R | ○ | ○ | ○ | ○ | ○ | ○ | | | | | | | |

***Note:*** *○ represents connectivity edges that are statistically significant following NBS correction at the specified primary threshold. Results are reported up to the first threshold at which no significant effects were observed; higher thresholds showed no significant effects and are omitted. DMN, default mode network; SN, salience network; CN, control network; DAN, dorsal attention network; L, left; R, right.*

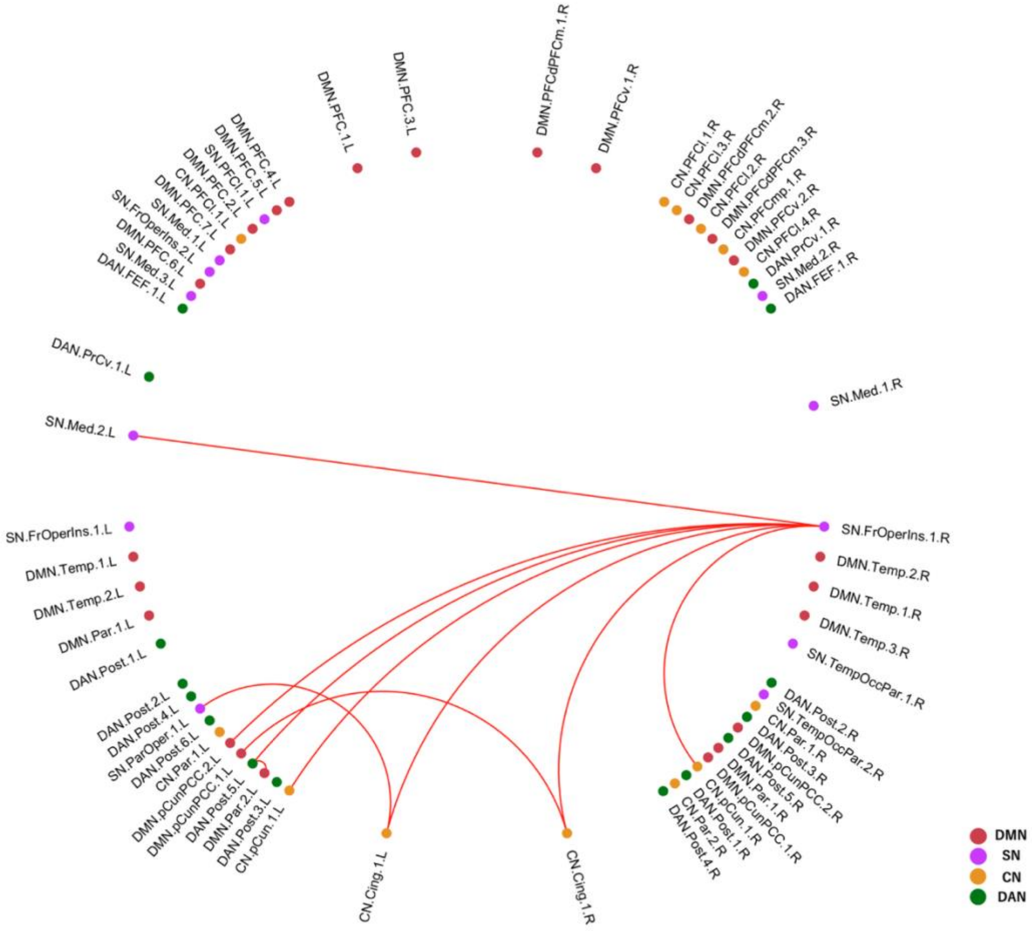


**Figure S9. F-statistic-based network-based statistic (NBS) results at $F_{primary}$ = 6.1 among deliberate MW, spontaneous MW, and on-task states at alpha band (7.5–12Hz).**

F-statistic-based NBS identified a significant main effect of cognitive state on functional connectivity in the alpha band (7.5–12 Hz; NBS-adjusted p < .0125, Bonferroni-corrected across four frequency bands). The significant network comprised 10 edges connecting 11 regions of interest, indicating differences in alpha band connectivity across deliberate MW, spontaneous MW, and on-task states.

***T-statistic NBS results between deliberate MW and spontaneous MW***

**Table S2. Alpha band (7.5–12Hz) NBS results (T-statistic) between deliberate mind-wandering and spontaneous mind-wandering across $T_{primary}$ thresholds from 2.0 to 3.5 (0.1 increments) NBS-adjusted $p < .0083$)**

| Edges | | $T_{primary}$ Threshold | | | | | | | | | | | | | | | |
|---|---|---|---|---|---|---|---|---|---|---|---|---|---|---|---|---|---|
| ROI 1 | ROI 2 | 2 | 2.1 | 2.2 | 2.3 | 2.4 | 2.5 | 2.6 | 2.7 | 2.8 | 2.9 | 3 | 3.1 | 3.2 | 3.3 | 3.4 | 3.5 |
| DMN.pCunPCC.1.L | CN.Cing.1.R | ○ | ○ | ○ | ○ | ○ | ○ | ○ | ○ | ○ | ○ | ○ | ○ | ○ | ○ | ○ | |
| SN.FrOperIns.1.R | CN.Cing.1.R | ○ | ○ | ○ | ○ | ○ | ○ | ○ | ○ | ○ | ○ | ○ | ○ | ○ | ○ | ○ | |
| SN.FrOperIns.1.R | CN.Cing.1.L | ○ | ○ | ○ | ○ | ○ | ○ | ○ | ○ | ○ | ○ | ○ | ○ | ○ | ○ | ○ | |
| SN.FrOperIns.1.R | CN.pCun.1.L | ○ | ○ | ○ | ○ | ○ | ○ | ○ | ○ | ○ | ○ | ○ | | | | | |
| SN.FrOperIns.1.R | CN.pCun.1.R | ○ | ○ | ○ | ○ | ○ | ○ | ○ | ○ | ○ | ○ | ○ | ○ | ○ | ○ | ○ | |
| SN.FrOperIns.1.R | DMN.pCunPCC.2.L | ○ | ○ | ○ | ○ | ○ | ○ | ○ | ○ | ○ | | | | | | | |
| SN.FrOperIns.1.R | SN.Med.2.L | ○ | ○ | ○ | ○ | ○ | ○ | | | | | | | | | | |

***Note:*** *○ represents connectivity edges that are statistically significant following NBS correction at the specified primary threshold. Results are reported up to the first threshold at which no significant effects were observed; higher thresholds showed no significant effects and are omitted. FrOperIns, frontal operculum-insula; Cing, cingulate; pCunPCC, precuneus posterior cingulate cortex; Med, medial; DMN, default mode network; SN, salience network; CN, control network; DAN, dorsal attention network; L, left; R, right.*

### *T-statistic NBS results between deliberate MW and on-task state*

**Table S3. Alpha band (7.5–12Hz) NBS results (T-statistic) between deliberate mind-wandering and on-task states across $T_{primary}$ thresholds from 2.0 to 3.9 (0.1 increments)**

| Edges | | $T_{primary}$ Threshold | | | | | | | | | | | | | | | | | | | |
|---|---|---|---|---|---|---|---|---|---|---|---|---|---|---|---|---|---|---|---|---|---|
| **ROI 1** | **ROI 2** | **2.0** | **2.1** | **2.2** | **2.3** | **2.4** | **2.5** | **2.6** | **2.7** | **2.8** | **2.9** | **3.0** | **3.1** | **3.2** | **3.3** | **3.4** | **3.5** | **3.6** | **3.7** | **3.8** | **3.9** |
| SN.FrOperIns.1.R | CN.Cing.1.L | ○ | ○ | ○ | | | | | | | | | | | | | | | | | |
| SN.FrOperIns.1.R | CN.Cing.1.R | ○ | ○ | ○ | ○ | ○ | ○ | ○ | | | | | | | | | | | | | |
| SN.FrOperIns.1.R | CN.pCun.1.L | ○ | ○ | ○ | ○ | ○ | ○ | ○ | ○ | ○ | ○ | ○ | ○ | ○ | ○ | ○ | | | | | |
| SN.FrOperIns.1.R | CN.pCun.1.R | ○ | ○ | ○ | ○ | ○ | ○ | ○ | ○ | ○ | ○ | ○ | ○ | | | | | | | | |
| SN.FrOperIns.1.R | SN.Med.2.L | ○ | ○ | ○ | ○ | ○ | ○ | ○ | ○ | ○ | ○ | ○ | ○ | ○ | ○ | ○ | ○ | | | | |
| SN.FrOperIns.1.R | DMN.pCunPCC.2.L | ○ | ○ | ○ | ○ | ○ | ○ | ○ | ○ | ○ | ○ | ○ | ○ | ○ | ○ | ○ | ○ | | | | |
| SN.FrOperIns.1.R | DAN.Post.5.L | ○ | ○ | ○ | ○ | ○ | ○ | ○ | ○ | ○ | ○ | | | | | | | | | | |
| DAN.Post.5.L | DMN.Par.2.L | ○ | ○ | ○ | ○ | ○ | ○ | ○ | ○ | ○ | ○ | | | | | | | ○ | ○ | ○ | |

| | | | | |
|---|---|---|---|---|
| SN.ParOper.1.L | CN.Cing.1.L | ○ | ○ | ○ |

***Note:*** *○ represents connectivity edges that are statistically significant following NBS correction at the specified primary threshold. Results are reported up to the first threshold at which no significant effects were observed; higher thresholds showed no significant effects and are omitted. FrOperIns, frontal operculum-insula; Cing, cingulate; pCunPCC, precuneus posterior cingulate cortex; Med, medial; Par, parietal; Post, Posterior; DMN, default mode network; SN, salience network; CN, control network; DAN, dorsal attention network; L, left; R, right.*
